# Use of RHO GTPase by L1 GABAergic neurons in frontal cortex and L6 glutamatergic neurons in prefrontal cortex differentiates states of unconsciousness

**DOI:** 10.1101/2025.06.21.660840

**Authors:** Y-H Taguchi, Turki Turki

**Affiliations:** Department of Physics, Chuo University, 1-13-27 Kasuga, Bunkyo-ku, Tokyo, 112-8551, Japan; Department of Computer Science, King Abdulaziz University, Jeddah, 21589, Saudi Arabia

**Author notes:** Contributing authors.

**Keywords:** anaesthesia, sleep, tensor decomposition, feature selection, AI consciousness studies

## Abstract

Unconsciousness can be induced by sleep and general anaesthesia alike; however, the extent to which these states differ is not well understood. In this study, we aimed to determine the similarities and differences between the unconscious states caused by sleep and those caused by anaesthesia, as well as investigate gene expression profiles by applying tensor decomposition-based unsupervised feature extraction to the two gene expression profiles. One of the two expression profiles was obtained from mice treated with sevoflurane, a type of inhaled anaesthesia, whereas the other two expression profiles were obtained from sleeping and awake mice. We selected two sets of genes (507 and 1048 genes) that were distinctly expressed between sleep- or anaesthesia-induced unconsciousness and the awake state. Both sets of genes include various distinct genes which are common to the state of unconsciousness during sleep and anaesthesia. The former was enriched in GABAergic synapses and the first layer of the frontal cortex, whereas the latter was enriched in glutamatergic synapses and the sixth layer of the prefrontal cortex. Additionally, both sets of genes were enriched in the RHO GTPase pathway. Based on these results, we hypothesised that L1 GABAergic neurons in the frontal cortex and L6 glutamatergic neurons in the prefrontal cortex use RHO GTPase to differentiate between the states of unconsciousness induced by general anaesthesia and sleep.

## 1 Introduction

Although sleep and general anaesthetics can both induce a state of unconsciousness, the resulting unconscious states are not necessarily equivalent [1, 2]. Unconscious but arousable sedation is closely related to sleep-wake circuits, whereas unconscious and unarousable anaesthesia are independent. General anaesthesia is notable for its ability to decrease sleep propensity. Conversely, increased sleep propensity due to insufficient sleep potentiates the anaesthetic effects [3].

Natural sleep is an endogenously generated state involving the active suppression of consciousness in specific brain regions, including the nuclei in the brainstem, diencephalon, and basal forebrain. The sleep cycle oscillates between the rapid eye movement (REM) and non-rapid eye movement (NREM) stages. In contrast, anaesthesia-induced unconsciousness is a pharmacologically induced and reversible state of unconsciousness characterised by hypnosis, analgesia, amnesia, and akinesia. Unconsciousness is achieved through the systemic administration of anaesthetics that disrupt neural communication, leading to a more widespread suppression of brain activity than that observed in natural sleep [4, 5].

Neurophysiologically, the brain exhibits slow-wave activity (SWA) with specific connectivity patterns during NREM sleep. By contrast, general anaesthesia often induces a state of burst suppression, indicating a profound disruption of thalamocortical connectivity and impaired information processing [4]. In addition, anaesthesia disrupts thalamocortical connectivity more than natural sleep, resulting in impaired information integration and processing [4, 6–8].

Unlike natural sleep, which is essential for physiological restoration and memory consolidation, anaesthesia-induced unconsciousness does not provide restorative benefits. Moreover, anaesthesia can suppress REM sleep, leading to a deficit that the body may attempt to recover post-anaesthesia.

In summary, although anaesthesia-induced unconsciousness and natural sleep may appear superficially similar, they differ significantly in their underlying mechanisms, neurophysiological characteristics, and functional outcomes.

This study aimed to understand the differences and similarities between sleep and anaesthesia from a genomic perspective. To this end, we applied the previously proposed tensor-decomposition-based unsupervised feature extraction to both sleep and anaesthesia gene expression profiles and compared them with those previously reported in [9]. We identified several hundred genes whose expression profiles were shared between the sleep and anaesthesia groups. These genes are involved in sleep- and anaesthesia-associated biological processes.

## 2 Results

Figure 1 provides an overview of the study design.

**Fig. 1.**
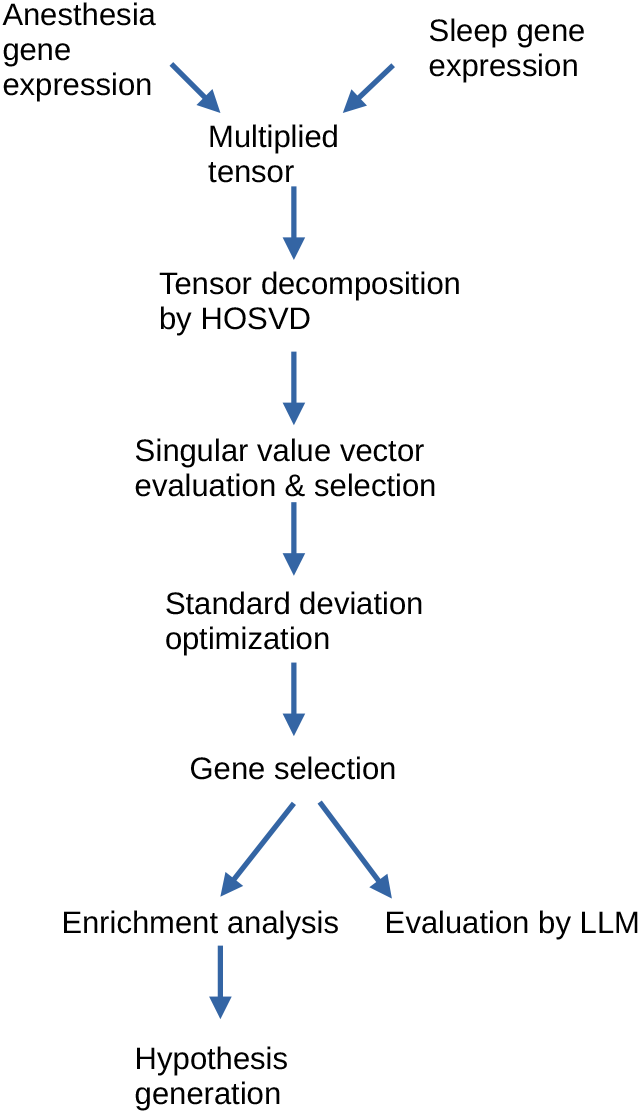
Schematic representation of the analyses performed in this study. First, two gene expression profiles were multiplied to generate a tensor. Tensor decomposition was applied to the generated tensor and singular value vectors were evaluated. After the selection of the singular value vectors attributed to various experimental conditions (e.g., tissue specificity and sleep vs. awake), the singular value vectors attributed to genes and used to select genes were selected. After optimizing the standard deviation, *P* -values were attributed to genes, and genes associated with adjusted *P* -values less than the threshold value (*<* 0.01) were selected. The selected genes were evaluated using large language model (LLM) and enrichment analysis, based on which we generated the hypothesis.

### 2.1 Gene selection

After applying HOSVD [10] to *x*_*ijkk*_*′*_*mm*_*′*, we obtained singular value vectors. Because we wanted to identify genes whose expression was distinct between sleep or anaesthesia and wake or control without any other dependencies, we selected the singular value vectors shown in Fig. 2 (Top left: *u*_2*j*_, top center: *u*_1*k*_, Top right: *u*_1*k*_*′*, bottom left and right: *u*_1*m*_, bottom centre: *u*_1*m*_*′*). We also investigated which *G*(*ℓ*_1_, 2, 1, 1, 1, 1) had the largest absolute value and selected *ℓ*_1_ = 3 (Fig. 2, bottom right).

**Fig. 2.**
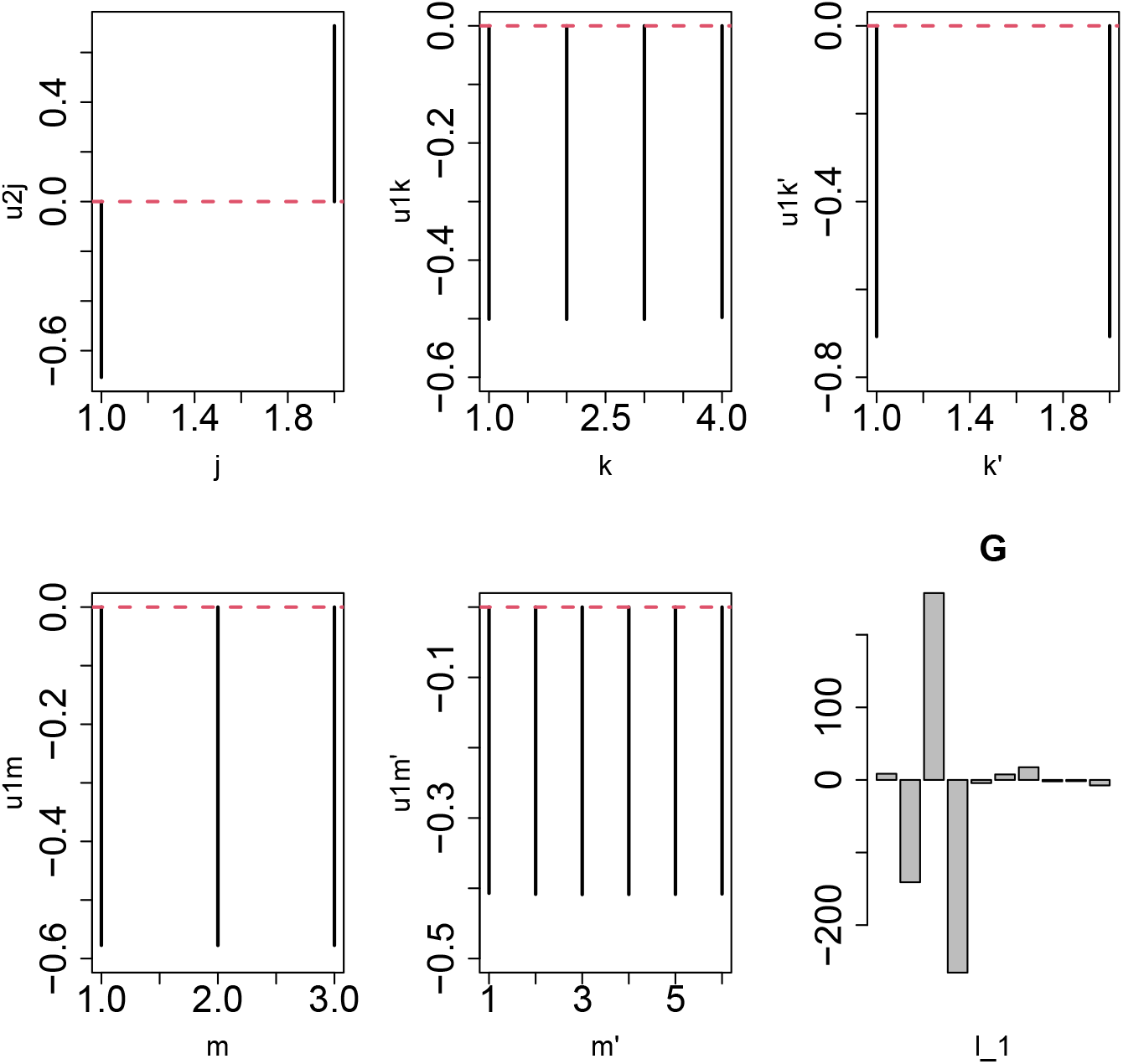
Singular value vectors and core tensor G. Top left: *u*_2*j*_, top center: *u*_1*k*_, top right: *u*_1*k*_*′*, bottom left and right: *u*_1*m*_, bottom center: *u*_1*m*_*′*, bottom right: *G*(*ℓ*_1_, 2, 1, 1, 1, 1)

Next, we attempted to optimise 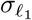, such that *P* -values computed using eq. (5) obeyed the null hypothesis that 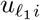 follows the Gaussian distribution as much as possible (Fig. 3, right panel). As expected, the standard deviation of the histogram of *P*_*i*_ had minimum values at a specific 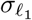 and the histogram of 1 − *P*_*i*_ followed a uniform distribution, indicating that the null hypothesis was as accurate as possible, excluding the peak near *P*_*i*_ = 0, which corresponded to the selected genes. In particular, 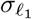, which had the minimum standard deviation in the histogram of *P*_*i*_, can be uniquely defined because only one minimum exists (Fig. 3, left panel). By setting the threshold value to 0.01, we selected 948 gene symbols associated with the adjusted *P* -values of *<* 0.01.

**Fig. 3.**
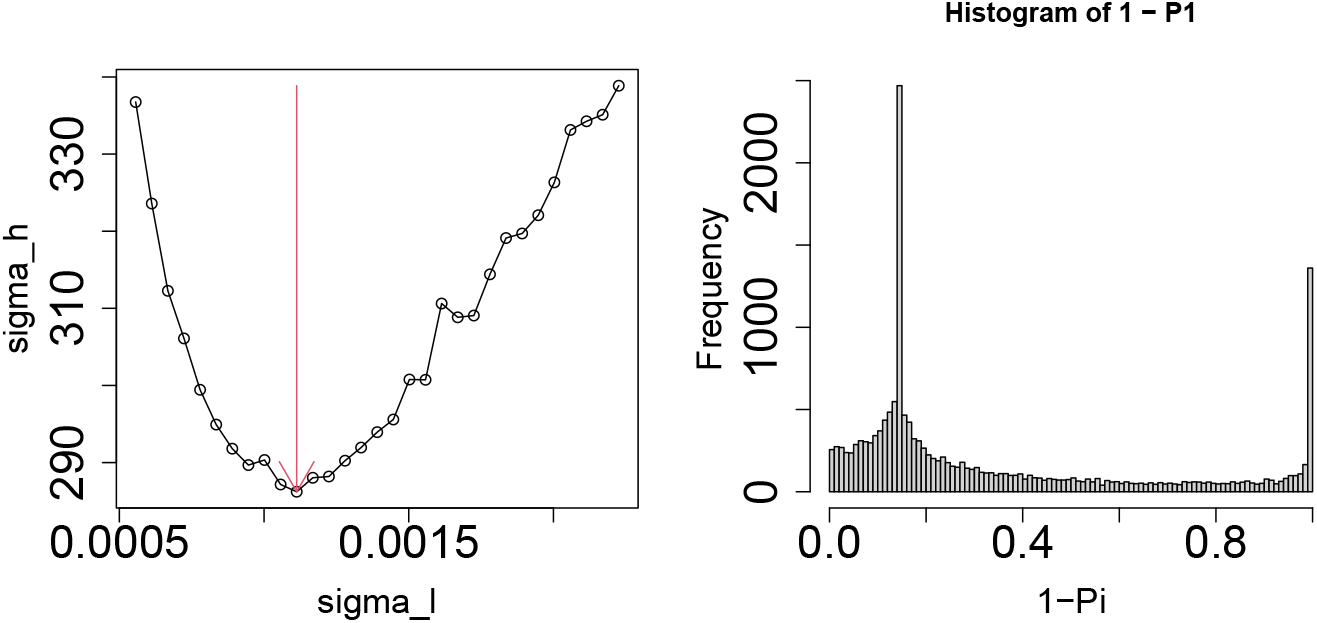
Left: standard deviation of histogram of *P*_*i*_ as a function of 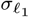 for *ℓ*_1_ = 3. The vertical red arrow suggests the selected value of 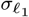. Right: histogram of 1 − *P*_*i*_ for the optimal 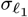.

Genes associated with 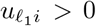 were distinct from those associated with 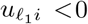, which were not associated with any enriched terms. In the following section, we consider only the 507 genes associated with 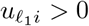.

We also consider *ℓ*_1_ = 4 is associated with the second-largest *G*. We attempted to optimise 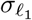 (Fig. 4, left panel).

**Fig. 4.**
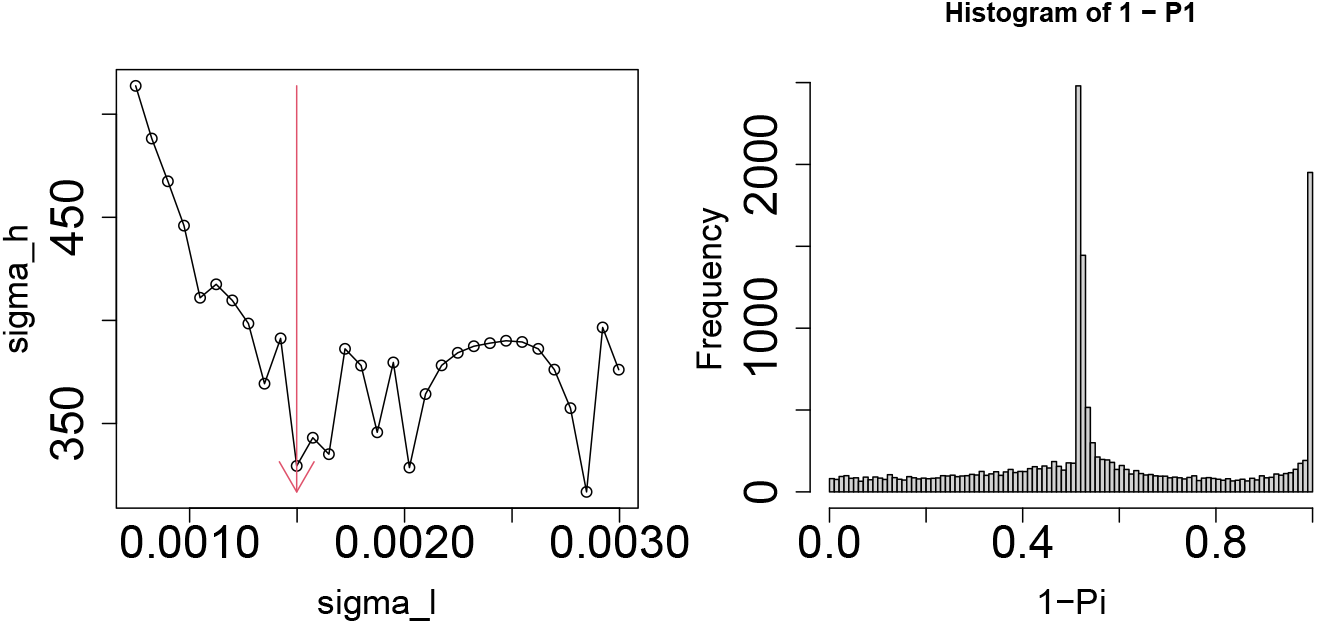
Left: standard deviation of histogram of *P*_*i*_ as a function of 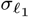 for *ℓ*_1_ = 4. The vertical red arrow suggests the selected value of 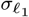. Right: histogram of 1 − *P*_*i*_ for the optimal 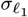.

In contrast to *ℓ*_1_ = 3, where 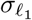 is associated with the smallest standard deviation, the histogram of *P*_*i*_ is unclear because many local minima have nearly the same standard deviation values. We decided to use the smallest possible 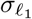, which resulted in the largest number of selected genes. It is clear that the Gaussian distribution is well optimized (Fig. 4, right panel). A total of 1,512 genes were selected. If we divide the set of genes by the sign of 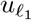, then only 1,048 genes associated with 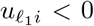 were enriched in various biological terms. Therefore, we considered these 1,048 genes.

*ℓ*_1_ = 3 genes with 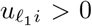 and *ℓ*_1_ = 4 genes with 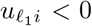 are valid because the sign of *G* is inverted between *ℓ*_1_ = 3 and *ℓ*_1_ = 4 (Fig. 2, bottom right). As can be seen in eq. (4), not individuals 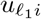 or *G* but their products 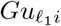 are important. Thus, for a *G* with the opposite sign, 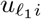 must have the opposite sign.

Figure 5 shows a Venn diagram of the 507 genes and 1,048 genes. Although there was some overlap, the majority of genes were distinct between the two sets. Thus, it is worthwhile to consider two sets of genes (for the intersection, we can hypothesise the origin later, see below).

**Fig. 5.**
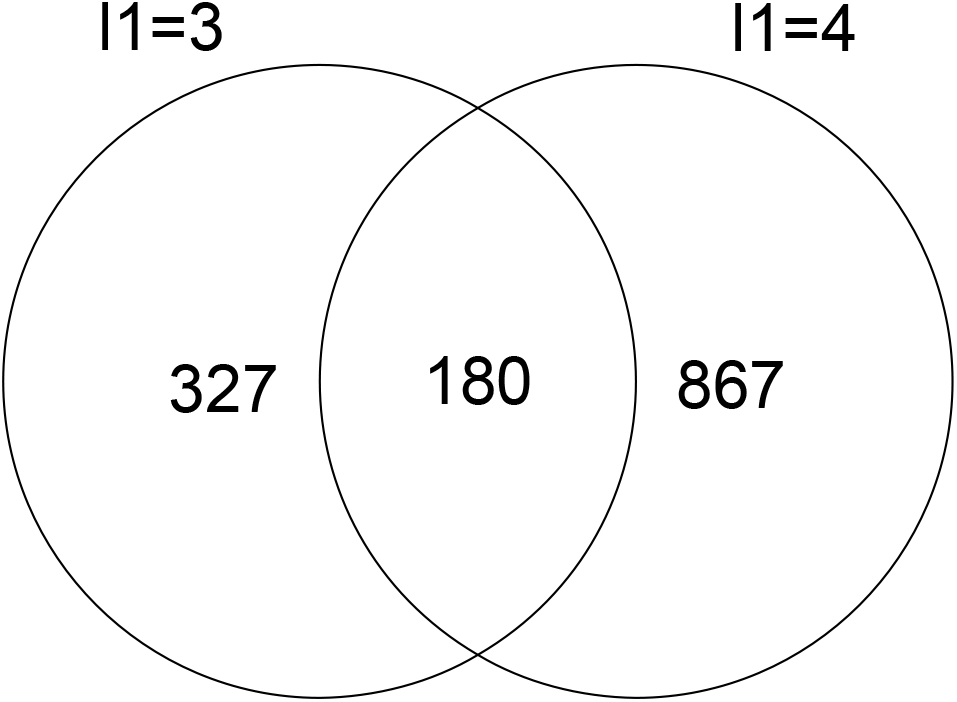
Left: standard deviation of histogram of *P*_*i*_ as a function of 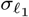 for *ℓ*_1_ = 4. The vertical red arrow suggests the selected value of 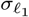. Right: histogram of 1 − *P*_*i*_ for the optimal 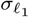.

### 2.2 Evaluation by LLM

First, we used the selected 507 and 1,048 genes separately in various large language models (LLM) to determine whether there were genes related to both sleep and anaesthesia. The prompt used was “This list of genes has been listed as being related to sleep and anaesthesia. Please review these lists, select those that you think are relevant to both sleep and anaesthesia, and summarise them with citations to supporting articles”, or its Japanese translation (Table 1). Several genes were identified in multiple LLMs, suggesting that the results are reliable.

**Table 1.**
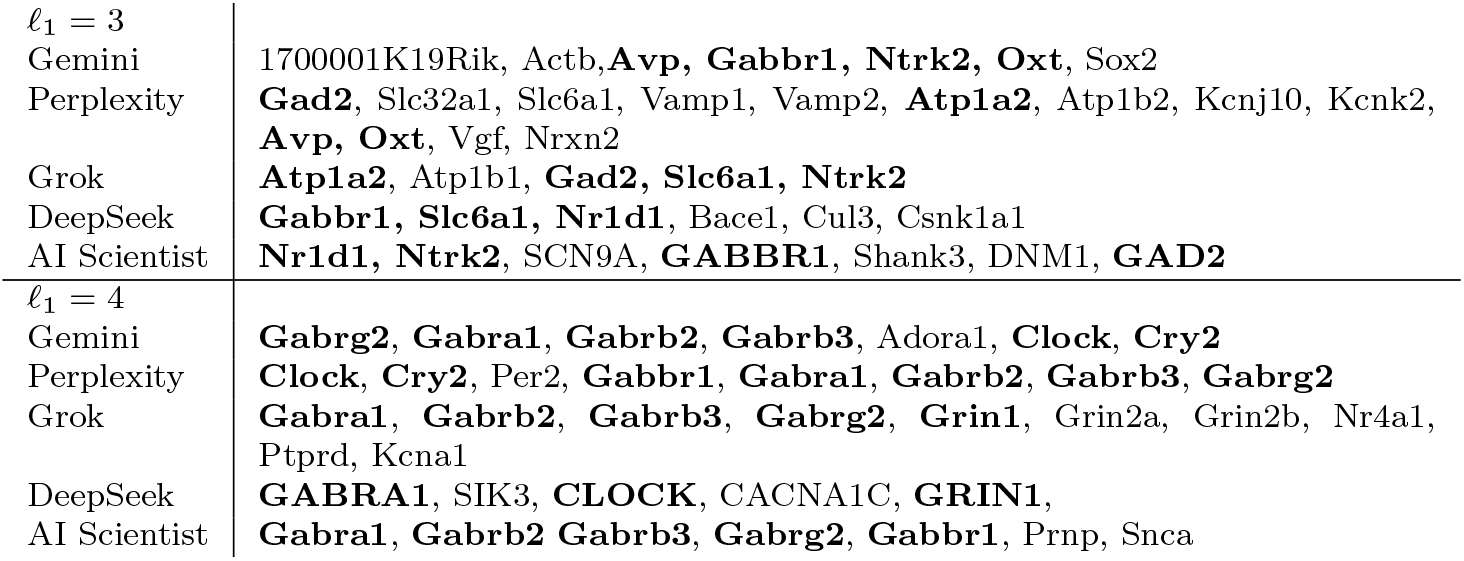
Sleep- and anaesthesia-related genes reported by various LLMs (bold).

Notably, the genes selected were relatively distinct between *ℓ*_1_ = 3 and *ℓ*_1_ = 4, indicating that considering *ℓ*_1_ = 3, 4 separately is informative despite the two sharing hundreds of genes(Fig. 5). In the following section, we summarise the references based on which the LLMs selected genes. This is to highlight the fact we did not simply trust the results of the LLMs but evaluated them based on the papers cited by the models. In addition, none of the genes selected by the LLMs were omitted, and all the genes included are listed in Table 1.

#### 2.2.1 AVP (*ℓ*_1_ = 3)

AVP is an important neuronal peptide in suprachiasmatic nucleus (SCN), acting as a master clock and controlling the circadian rhythm, a key mechanism in wake-sleep regulation. The AVP signal controls SCN network characteristics. AVP neurons in the SCN exhibit rhythmic calcium activity [11]. The arginine vasopressin neurons in the paraventricular nucleus of the hypothalamus (PVH^AVP^) promote arousal [12]. AVP participates in REM sleep regulation under normal physiological conditions [13]. It also plays an important role in maintaining circulatory stability, especially in situations in which the sympathetic and renin-angiotensin systems are suppressed (e.g. during anaesthesia or in shock) [14].

#### 2.2.2 Gabbr1(*ℓ*_1_ = 3, 4), Gabra1, Gabrb2, Gabrb3, and Gabrg2 (*ℓ*_1_ = 4)

GABA B receptor signalling is involved in sleep regulation, particularly slow-wave sleep (SWS). GABA B agonists, such as baclofen and *γ*-hydroxybutyrate (GHB)/sodium oxybate, increase SWS and delta wave power in animals and humans, with minimal effects on REM sleep. By contrast, GABA B antagonists reduce SWS. In preclinical studies, GABA B agonists have been associated with the increased expression of genes related to SWS production and circadian function [15].

The GABAergic pathway is central to the action of many general anaesthetics (e.g. propofol, isoflurane, sevoflurane, etomidate, and barbiturates), which often target GABA A receptors [16]. GABA B receptors may also be involved, and the GABA B agonist, baclofen, has shown potential therapeutic effects in animal models of alcohol dependence (reduced self-administration behaviour) [17]. In a study using GABAB flox mice and astrocyte-specific ablation using a viral vector (AAV-Cre) during surgery under isoflurane anaesthesia, Gabbr1 was ablated, demonstrating the utility of these models for in vivo GABA B function studies. This technique avoids the developmental problems associated with constitutive knockout (KO) [18]. Although many anaesthetic agents act primarily on GABA A receptors, GABA B receptors modulate the overall network inhibition and may contribute to anaesthetic states, particularly specific elements such as sedation and muscle relaxation, as suggested by their functions and agonist effects. However, GABA B receptors are the primary targets of common anaesthetic agents, such as propofol and isoflurane, and are not as prominent in these snippets as the GABA A receptor. However, there is evidence for the direct involvement of the GABA A receptor in these snippets.

Gabbr1 is a member of the gamma-aminobutyric acid (GABA) receptor gene family, which encodes receptors for GABA, the principal inhibitory neurotransmitter in the central nervous system. GABAergic neurotransmission is a well-established mediator of sleep induction and of the sedative effects of many anaesthetic agents, including propofol. Variants of GABA receptor genes have been associated with altered anaesthetic sensitivity and modulation of sleep-promoting circuits, underscoring the dual role of Gabbr1 in these processes [2, 19].

#### 2.2.3 Ntrk2 (*ℓ*_1_ = 3)

In the brain of rats, Ntrk2 shows diurnal variations in expression levels and the use of selective polyadenylation (APA) sites. Specific APA isoforms are altered by sleep deprivation and subsequent restorative sleep [20], suggesting that Ntrk2 regulation at the mRNA processing level is associated with circadian and sleep homeostasis mechanisms. Ntrk2 has been identified as one of 46 genes in which circadian-related APAs overlap with those associated with genes associated with susceptibility to brain disease [20]. In studies using yak testes, lncRNAs (MSTRG.19083.1) were identified as miR-429-y and were shown to function as sponges for Ntrk2 to regulate circadian rhythms via the cAMP pathway, suggesting the existence of a complex regulatory network in circadian processes involving Ntrk2 (even outside the brain) [21].

Genetic polymorphisms in human NTRK2 are associated with symptom burden in patients with breast cancer undergoing chemotherapy. Specifically, the T allele of SNP rs1212171 was associated with increased sleep disruption and fatigue interference during treatment [22]. This suggests that individual genetic variants in the TrkB pathway may influence susceptibility to common treatment side effects, such as sleep interruption and fatigue, suggesting that they may affect susceptibility to common treatment side effects, such as fatigue, possibly via alterations in neurotrophic factor signalling and interactions with inflammatory responses [22]. Based on the data provided, Ntrk2 is directly involved in the anaesthetic mechanism (e.g. LORR). Although there is no evidence, its role in neuroplasticity, survival, and the regulation of the circadian rhythm suggests that it may be a modulator of anaesthetic sensitivity and recovery. The most direct link is its association [22] with sleep disorders.

Ntrk2 encodes the neurotrophic receptor tyrosine kinase B (TrkB), which is involved in neuroplasticity, synaptic regulation, and learning. Ntrk2 expression is modulated following isoflurane anaesthesia, indicating that Ntrk2 may influence the neural circuits that underlie both sleep and anaesthetic states. Given that proper synaptic function and plasticity are essential for maintaining stable sleep patterns and for the emergence of anaesthetic-induced unconsciousness, Ntrk2 has emerged as an important candidate in the shared biology of these processes [23].

#### 2.2.4 Oxt (*ℓ*_1_ = 3)

Oxt signalling via the paraventricular nucleus (PVN) to the prefrontal cortex (PrL) pathway modulates the inhibitory balance of PrL (PV/VIP interneurons) during REM sleep and supports social memory fixation. OXT release occurs during REM sleep, wherein a lack of OXT signalling (e.g. due to chronic sleep deprivation) disrupts this balance and leads to social memory impairment. The activation of the REM sleepspecific PVN^OCT^ → PrL pathway or intranasal OXT administration rescues these deficits in sleep-deprived mice [24]. This suggests that it does not act universally on social memory, but regulates specific cortical circuits during a particular offline brain state (REM sleep), which is important for memory consolidation. Polymorphisms in the human OXTR gene (rs2254298, A allele) are associated with symptoms of obstructive sleep apnoea (OSA), particularly shortness of breath, snoring, and reduced subjective sleep efficiency [25]. This suggests that endogenous OXT signalling may be involved in breathing regulation during sleep. OXT levels, social support, and sleep quality in HIV-positive women, with some studies demonstrating an association.

OXT reduces ethanol consumption and motivation in mice. However, this is not due to a general sedative effect [26]. Oxtr-KO mice do not exhibit ethanol-induced conditioned social preferences [27]. OXT administration (intranasal or intraperitoneal) is a potential treatment for ASD-related behaviours in animal models (VPA rats) to study its effects. However, this usually involves behavioural testing under baseline or manipulated conditions, rather than under anaesthesia [28]. These findings suggest a clinical context in which OXT-related systems may interact with anaesthetics; however, the interaction itself was not investigated.

#### 2.2.5 GAD2 (*ℓ*_1_ = 3)

GAD2 is a promising candidate for triggering conscious mechanisms of action. This gene encodes glutamate decarboxylase 2, an enzyme that is crucial for the synthesis of GABA, a primary inhibitory neurotransmitter in the central nervous system. As both natural sleep and various anaesthetic agents rely on the enhancement of inhibitory neurotransmission to induce loss of consciousness, GAD2 is functionally important in these processes. Increased GABA synthesis due to GAD2 activity can modulate the inhibitory tone in key sleep-regulating circuits and contribute to the depth of anaesthetic states [2, 29].

GAD2^LPO^ neuronal stimulation did not merely trigger wakefulness; the awake state produced by this stimulation was characterised by increased EEG theta activity. In contrast, the unilateral inhibition of GAD2^LPO^ neurons decreased the drive for arousal, as reflected by the persistent increase in NREM EEG SWA throughout the day. In summary, our experiments demonstrate the important role of GAD2^LPO^ neurons not only in the control of state transitions but also in linking arousal to sleep homeostasis [30].

#### 2.2.6 Alpha2 (*ℓ*_1_ = 3)

Evidence for Atp1a2 is more indirect. This gene encodes the catalytic and regulatory subunits of the sodium potassium pump. This pump maintains the ionic gradients across the cell membrane and is essential for neuronal excitability and transmission. Alpha2 agonist anaesthetics hyperpolarise neurons and inhibit action potential generation, for example, by activating inwardly rectifying potassium channels [31]. Alpha2 is also involved in ion regulation and contributes to sleep and anaesthesia regulation.

The fact that REM sleep deprivation increases Na-KATPase activity highlights the importance of ion balance in sleep regulation [32]. Under general anaesthesia, alpha2 may be responsible for altered neural activity under anaesthetic conditions [33].

#### 2.2.7 Slc6a1 (*ℓ*_1_ = 3)

The evidence for Slc6a1 is also indirect. Slc6a1 encodes GABA transporter 1 (GAT-1), which reuptakes GABA from synapses and regulates its levels. Anaesthetic drugs exert neuroinhibitory effects via GABA transporters. Dysfunction of the GABA system has been observed in models of sleep disorders, and mutations in this transporter have been found to affect sleep regulation [34].

#### 2.2.8 Nr1d1 (*ℓ*_1_ = 3)

Nr1d1 encodes a nuclear receptor that is a key transcriptional regulator of the circadian clock machinery. Its expression and central position in gene networks are significantly affected by isoflurane anaesthesia, suggesting that changes in Nr1d1 levels can alter circadian timing, mood, and memory. Its dual role in maintaining normal sleep–wake cycles and modulating anaesthetic sensitivity supports its relevance as a molecular bridge between sleep and the regulation of anaesthesia [23].

#### 2.2.9 Clock (*ℓ*_1_ = 4)

Clock genes are fundamental in daily sleep timing and circadian rhythm regulation [31]. General anaesthesia using isoflurane, sevoflurane, propofol, and dexmedetomidine may disrupt circadian rhythms and alter clock gene expression (e.g. Per2 and Cry) in the master clock [35].

#### 2.2.10 Cry2 (*ℓ*_1_ = 4)

Cry2 mutations are associated with familial advanced sleep phase syndrome (FASP), which is characterised by early sleep and wake times, highlight the role of Cry2 in sleep-wake timing in humans [36]. General anaesthetics, such as isoflurane, alter clock gene expression, such as Cry-m in honeybees and Per2, which has often been studied in conjunction with Cry. This indicates that isoflurane delays the Cry-m mRNA oscillations [35].

### 2.3 Enrichr

By uploading the 507 selected genes for *ℓ*_1_ = 3 and 1,048 genes for *ℓ*_1_ = 4 to Enrichr, several enriched terms were identified.

#### 2.3.1 Reactome Pathways 2024

##### GABA synthesis, release, re-uptake, and degradation

GABA synthesis plays a critical role in consciousness [37–39]; it is the third highest ranked term in this category for *ℓ*_1_ = 3.

##### RHO GTPases

“RHO GTPases Activate Formins” and “RHO GTPase Effectors” are the sixth and seventh terms in this category for *ℓ*_1_ = 3, respectively, while “Signalling by Rho GTPases” and “Signalling by Rho GTPases, Miro GTPases and RHOBTB3” are the fifth and seventh terms in this category for *ℓ*_1_ = 4, respectively. Recently, the RHO GTPases pathway was shown to control sleep-wake rhythm. For example, SIK3, a kinase that controls mammalian sleep demand, can target a small RhoA protein, whose activity regulates sleep-wake controls [40]. Similarly, some general anaesthetics can bind to Rho GTPase regulators and affect Rho GTPase. Inhaled anaesthetics, such as halothane, can bind to RhoGDI and alter its functionality [41]. These results suggested that RHO GTPases are involved in consciousness.

##### Relationship with sleep

Rho family small GTPases (RhoA, Rac1, and Cdc42) act as molecular switches that govern synaptic structure and function in the brain and are deeply involved in plastic changes in neuronal circuits during sleep and wakefulness. During wakefulness, these pathways contribute to synaptic strengthening and maintain neuronal excitability in response to experiences, whereas during sleep, they play a dynamic role in promoting synapse reduction and reorganisation to stabilise brain networks. Animal model studies have shown that disruption of the Rho GTPase pathway leads to reduced sleep duration, abnormal sleep homeostasis, and cognitive dysfunction [42, 43], suggesting its involvement in human sleep, development, and neurological disorders.

With regards to anaesthesia, phenomena at the cellular level induced by anaesthetic drugs such as isoflurane, sevoflurane, and propofol cause neuronal dendritic spine loss and growth cone collapse, delayed astrocyte maturation, altered vascular endothelial barrier, and accelerated endothelial vesicle release, with small GTPase activity fluctuations in the Rho family. In particular, RhoA and its downstream Rho-kinase (ROCK) are hub molecules involved in anaesthetic-induced cytoskeletal changes, indicating that the manipulation of the RhoA/ROCK pathway may mitigate the adverse effects of anaesthetic drugs [44, 45].

#### 2.3.2 KEGG 2021 Human

##### GABAergic synapse

GABAergic synapses play a critical role in consciousness [37–39] and are the highest-ranked terms in this category for *ℓ*_1_ = 3 and the fourth top-ranked term for *ℓ*_1_ = 4.

##### Glutamatergic synapse

In addition, the glutamatergic synapse plays a critical role in consciousness [46] and is the highest-ranked term in this category for *ℓ*_1_ = 4.

##### Dopaminergic synapse

Dopaminergic synapses are related to both sleep and anaesthesia [47] and are the second top-ranked terms in this category for *ℓ*_1_ = 4.

##### Circadian entrainment

Circadian rhythms are known to play critical roles in sleep [48] and anaesthesia [49], and are the eighth top-ranked terms in this category for *ℓ*_1_ = 4.

##### Thyroid hormone signalling pathway

The “thyroid hormone signalling pathway” is the third highest ranked term in this category for *ℓ*_1_ = 3. The thyroid gland produces two primary hormones [50].

- Thyroxine (T4): A prohormone.
- Triiodothyronine (T3): The active form.

T4 is converted to T3 by deiodinases (especially type 2 Deiodinase (DIO2) in the brain. T3 binds to the thyroid hormone receptors (TR*α* and TR*β*), which are nuclear receptors that regulate gene expression. These receptors are highly expressed in brain regions critical for consciousness, including the cortex, thalamus, and reticular formation of the brainstem [51]. Neurophysiologically, the thyroid hormone signalling pathway affects consciousness. T3 increases neuronal excitability by regulating Na^+^/K^+^ ATPase expression (maintaining membrane potential) [52] and ion channel function (e.g. *K*^+^ channels) [53]. T3 also enhances synaptic plasticity in areas like the prefrontal cortex [54] and hippocampus [55], supporting higher-order functions tied to consciousness (e.g., attention, memory, and awareness). Thalamocortical circuits regulate arousal states and sleep-wake transitions [56–58]. Thyroid hormones modulate the neurotransmitter systems (e.g. glutamate, GABA, and acetylcholine) involved in thalamocortical oscillations that underlie different states of consciousness [59, 60]. These observations strongly suggest that the thyroid hormone signalling pathway regulates consciousness, although the thyroid hormone signalling pathway is not likely to play a critical role.

#### 2.3.3 Allen Brain Atlas down

This category lists genes whose expression was lower in the treatment group than in the control group. Since the control and treatments vary between experiments, the distinction between “Allen Brain Atlas down” and “Allen Brain Atlas up” is not critical. Several important regions in this category have been identified during sleep. The top-ranked regions not listed here were also expected to be important during sleep.

##### Hindbrain

Hindbrain neurons flip the switch among REM sleep, non-REM sleep, and wakefulness [61]. The hind brain plays a critical role in anaesthesia [62]. “Hindbrain” is the top ranked term for *ℓ*_1_ = 4.

##### layer 1 of FCx

“Layer 1 of FCx” is the top ranked term for *ℓ*_1_ = 3. Layer 1 (L1) is also known as the bimolecular layer, and there are few neurons in L1. Instead, dendrites and axon terminals accumulate in L1. In particular, the apical tufts of pyramidal neurons were concentrated. There are some interneurons in Layer 1, which is GABAergic, such as neurogliaform cells. L1 is a hub of synaptic connections, such as corticocortical and thalamic inputs. L1 is the control centre of the pyramidal neurons in the lower layer based on incoming information. Attention control and conscious judgment are controlled by L1 of the frontal cortex (Fcx). For example, incoming signals from fronto-parietal network affect the activity of pyramidal neuron in layer 5 (L5). NMDA receptor-dependent synaptic plasticity is thought to be related to L1 of the FCx [63].

As mentioned above, L1 governs wide-ranging cortical regulation, especially topdown control, and changes in the state of consciousness. Regarding the relationship with sleep, L1 of the FCx is the source of delta brain waves, whose occurrence and propagation are important in slow-wave sleep (SWS) [64–66]. In contrast, general anaesthesia strongly suppressed corticocortical inputs and dendritic spikes. Dendritic spikes in pyramidal neurons and activity of lower-layer neurons were reduced. This results in suppression of global consciousness. For example, isoflurane and other anaesthetics suppress the coupling of L1 dendrites and somas in cortical pyramidal neurons [67]. This process is related to the unconsciousness. Ketamine affects L1 of the FCx differently from that of the deep layers [68]. In short, during deep sleep or anaesthesia, the neuromodulatory tone drops and L1 inputs diminish, which, along with strong L1 inhibition, contribute to the hyperpolarization of apical dendrites and synchronisation of slow oscillations.

##### FCx layer 2

“Layer 2 of FCx” is the second highest ranked term for *ℓ*_1_ = 3. In contrast to L1 which included only a few cell bodies, Layer 2 (L2) mainly included the cell bodies. Thus, L2 was expected to collaborate with L1. L2 receives thalamic inputs through L1, especially from higher-order thalamic nuclei, acts as a relay hub like L1, and plays a role in associative learning, memory encoding, and modulation of cognitive function. Neurons in L2 tend to have high input resistance and low firing thresholds, making them sensitive to synaptic inputs from L1 and other superficial layers [69]. L2 of the frontal cortex contains pyramidal cells and GABAergic interneurons, the latter of which are crucial for local circuit processing in the frontal cortex.

In NREM, especially in SWS, small pyramidal neurons and astrocytes (also known as interneurons which are missing in L1) in L2 of FCx exhibit states of large up- and downregulation, and their occurrence and propagation through L3 and L5 generate slow waves. Thalamocortical inputs toward L4 in FCx propagated with the help of L2 and L3 (L3 of FCx is the fifth top-ranked term) of FCx and generated slow-wave activity and sleep spindles in NREM, the latter of which play critical roles in memory consolidation in L2/L3 of the FCx [70].

General anaesthesia drastically altered the frequency of synaptic firing and synchronisation patterns at L2/L3. In particular, slow oscillation (0.1-1 Hz), which is analogous to the slow-wave activity in NREM, was enhanced. Nevertheless, the propagation speed and interaction between the layers can differ from SWS. Slow-wave propagation under anaesthesia from L1 to L5 through L2/L3 is relatively slower and more unstable than that during sleep [71]. Pyramidal cells in the L2/L3 phase are affected by the anaesthetic-induced overactivation of GABA receptors, inhibition of excitatory synaptic transmission, and intermittent firing. This causes a large-scale disruption of cortical networks, which leads to a loss of consciousness [72].

##### Primary motor area, Layer 1/Secondary motor area/Secondary motor area, layer 2/3

These are the eighth to tenth top-ranked terms in this category for *ℓ*_1_ = 3. In contrast to L1–L3 of FCx, which are important for NREM, the motor area is important for REM. In NREM, the primary motor cortex (M1) and premotor areas show reduced activity compared with wakefulness [73]. However, sleep spindles and slow oscillations in motor areas contribute to motor memory consolidation (e.g. learning new motor skills) [74]. During REM, the primary motor and premotor cortices are reactivated and often show activity levels similar to those observed during wakefulness [73]. Despite this cortical reactivation, motor output is suppressed by the brainstem (specifically the pontine reticular formation), which induces muscle atonia (paralysis) to prevent dreams [75]. This paradox (active motor areas but paralyzed muscles) is critical for dreaming without movement.

With regard to anaesthesia, motor cortex activity is strongly suppressed, but is not identical to sleep [76]. Similar to NREM sleep, motor output is abolished, but the cortical dynamics differ depending on the anaesthetic agent; propofol, sevoflurane, and isoflurane tend to produce strong, slow oscillations (0.1–1 Hz) in the motor cortex [77–79], reflecting deep cortical suppression. Some anaesthetics (e.g. ketamine) show preserved or increased gamma activity (30–80 Hz), even in motor areas, despite loss of consciousness [80]. In terms of motor evoked potentials (MEPs), transcranial magnetic stimulation-evoked (TMS) responses from the motor cortex are markedly reduced under anaesthesia [81]. This indicates that the motor output pathways (cortex to spinal cord) are functionally disconnected.

However, unlike sleep, anaesthesia does not consolidate motor memory. Studies comparing motor sequence learning before and after anaesthesia have shown no performance gains (opposite to sleep effects) [82]. This is likely because anaesthesia globally suppresses plasticity mechanisms in motor circuits.

##### Supraoptic nucleus/episupraoptic nucleus

Although “Supraoptic nucleus” and “episupraoptic nucleus” are not highly ranked terms but only 145th and 185th ranked terms with significant adjusted *P* -values (6.28 × 10^−4^ and 2.80 × 10^−4^, respectively) for *ℓ*_1_ = 3, a recent study found a novel group of GABAergic and glutamatergic neurons in the supraoptic nucleus (SON) of the hypothalamus that was activated by multiple anaesthetics and that promoted NREM sleep [38] and termed them anaesthesia-activated neurons (AAN). Notably, AAN was discovered by investigating AVP and OXT expression, which was also identified using LLM (Table 1). Although they were also associated with significant *P* -values for *ℓ*_1_ = 4, they were not high (as low as 42th and 72th, respectively).

#### 2.3.4 Allen Brain Atlas up

This category lists the genes whose expression increased in the treatment groups compared to the control group. Since the control and treatments vary between experiments, the distinction between “Allen Brain Atlas down” and “Allen Brain Atlas up” is not critical.

##### Layer 6 of PCx

The multiple elements of layer 6 of the PCX-related subregions were ranked. “Sublayer 6b of PCx” (2nd), “layer 6 of PCx” (3rd), and “sublayer 6a of PCx” (4th) for *ℓ*_1_ = 4. It is heterogeneous and is composed of various types of neurons, mostly glutamate [83]: corticothalamic (CT) pyramidal (glutamate), corticocortical (CC), pyramidal (glutamate), L6b/Drd1a-Cre+ pyramidal (glutamate), orexin-sensitive L6 pyramidal (glutamate), and GABAergic interneurons (GABA).

Cortical layer 6b mediates state-dependent changes in brain activity and the effects of orexin on waking and sleeping [84]. The presence of orexin-sensitive neurons, almost exclusively within the mPFC L6 [85], adds another layer of complexity and significance. Orexin is a neuropeptide critical for maintaining arousal and wakefulness. Anaesthetics suppress arousal [86]. Thus, interference by anaesthesia with orexin signalling or direct effects on these orexin-sensitive L6 neurons could represent a specific mechanism by which anaesthesia dampens arousal-related modulation of PFC activity at this laminar level, contributing to the hypnotic component of anaesthesia.

In contrast, general anaesthesia affects layer 6 of PCx in various ways.

- Propofol: This widely used GABAergic anaesthesia has marked effects on deep PFC neurons. In layer 5 of the medial PFC (mPFC), which shares functional characteristics and connectivity relevant to L6, propofol predominantly inhibits the firing of regular-spiking (RS) neurons (putative pyramidal cells) and causes a significant suppression of fast-spiking (FS) interneurons. While the majority of RS neurons are inhibited, a small subset may exhibit excitation [87]. Propofol also leads to a general reduction in neuronal excitability, an effect that can be particularly pronounced in the lateral PFC compared to sensory areas like the auditory cortex [88]. In in vitro cortical slice preparations, a clinically relevant concentration of propofol (3*µ*M) has been shown to reduce muscarine-induced persistent potentials and associated spiking activity [89].
- Sevoflurane: Another commonly used inhaled GABAergic anaesthetic, sevoflurane, exerts effects on mPFC L5 neurons that are largely similar to those of propofol. Sevoflurane primarily inhibits RS neurons and suppresses FS interneuron activity [87]. Compared to isoflurane, sevoflurane tends to produce a uniform suppression of single-unit activity across the cortex. This potent suppression may be related to its effects on glutamatergic transmission; sevoflurane more effectively reduces calcium-dependent glutamate release than isoflurane, especially at higher concentrations [90].
- Ketamine (NMDA antagonist): In stark contrast to GABAergic anaesthetics, ketamine induces predominantly excitatory effects on both RS and FS neurons in mPFC L5, although a subpopulation of neurons can still be inhibited [87]. A striking effect of ketamine is its ability to cause a rapid and widespread switch in spontaneously active pyramidal neuron populations across all cortical layers, including deep layers L5 (and by extension, L6), in various cortical regions (somatosensory, visual, motor, and retrosplenial cortices). This phenomenon involves the suppression of neurons that are normally active during wakefulness and the concurrent activation of previously silent neurons [91]. This profound reorganization of active neuronal ensembles occurs with both systemic administration and local cortical application of ketamine, suggesting a fundamental alteration in cortical state rather than simple suppression. At low, therapeutically relevant concentrations (e.g., 1*µ*M) in mPFC L5 pyramidal neurons, ketamine has been shown to decrease the frequency of spontaneous inhibitory postsynaptic currents (sIPSCs) while increasing the frequency and amplitude of spontaneous excitatory postsynaptic currents (sEPSCs). This is consistent with a preferential blockade of NMDARs located on GABAergic interneurons, leading to pyramidal cell disinhibition. High concentrations of ketamine (e.g., 10 µM) tend to decrease the rates of both sIPSCs and sEPSCs [92]. In cortical slices, 20*µ*M ketamine appeared to enhance muscarine-induced persistent potentials and spiking (though non-significantly), whereas a much higher dose (100*µ*M) suppressed them [89]. This dichotomy between the general suppressive effects of GABAergic anaesthetics and the complex, often excitatory and state-switching effects of ketamine on L6/deep layer neurons suggests fundamentally different mechanisms by which these agents disrupt L6 processing and, consequently, consciousness.
- Isoflurane: This inhaled anaesthetic also modulates activity in deep cortical layers, including PFC infragranular layers, in a complex, frequency-specific manner (detailed in section 3.2). Compared to sevoflurane, isoflurane causes a less uniform suppression of neuronal firing [90].

Regarding firing patterns, the anaesthetics differentially affected RS and FS neurons. For GABAergic agents, such as sevoflurane and propofol, the activity of inhibited RS neurons in mPFC L5 is closely correlated with the transitions between loss of righting reflex (LORR) and recovery of righting reflex (RORR), highlighting their potential role in tracking anaesthetic depth. FS interneurons are generally significantly suppressed under these GABAergic anaesthetics. However, ketamine has mixed (inhibitory and excitatory) effects on FS neurons. Anaesthetics also influence bursting activity. For example, urethane can increase burst duration and the number of spikes within a burst in cortical neurons. The arousal-promoting peptide orexin, which acts on specific L6 neurons, increases burst frequency in the mPFC even under isoflurane anaesthesia, suggesting that some intrinsic circuit dynamics can be modulated regardless of the anaesthesia state.

#### 2.3.5 Validation using single cell data bases

Although many cellular levels of enrichment were identified, we used tissue-level measurements. To fill this gap, we also checked for enrichment in single cell-based categories. As can be seen below, the enrichment for two single-cell-based databases, “Azimth cell types 20021” and “Azimuth 2023” in Enrichr, is at least partially consistent with the above. Thus, our results may also be effective in single-cell-based analysis.

##### Azimuth cell types 2021

“SST+ DEFB108B+ Layer 1 GABAergic Neuron CL0000617” is the sixth highest ranked for *ℓ*_1_ = 3 and “Glutamatergic Neuron CL0000679” is top-ranked for *ℓ*_1_ = 4.

##### Azimuth 2023

“Fetal Development-L1-Limbic System Neurons” and “Fetal Development-L2-Limbic System Neurons” are top and the second ranked for *ℓ*_1_ = 3. “Mouse Motor Cortexcross-species cluster-Layer 6 Glutamatergic Neuron, Intratelencephalon-Projecting 2” is the sixth ranked for *ℓ*_1_ = 4.

## 3 Methods

### 3.1 Gene expression profile

#### 3.1.1 Gene expression profiles under anaesthesia treatment

Gene expression profiles under anaesthesia were generated using DRA010292 [9]. Three mice were treated with sevoflurane and three mice were taken as controls; in total, six mice were investigated. However, because this dataset was taken from the above DRA010292, we did not perform any experiments ourselves. The Fastq files of RNA expression in brain sections (hippocampus, hypothalamus, medial prefrontal cortex, and striatum) were generated. The files were mapped onto the mouse genome (mm10) using bowtie2 [93]. bowtie2 was used over of other transcriptome mappers because the gene expression profiles to be compared were measured using a DNA array rather than a transcriptome array (see the subsection below). Therefore, mapping short reads to the genome using not a transcriptome mapper but a DNA mapper is preferable. In total, we had two conditions (i.e. control and anaesthesia groups) and three mice for four brain subsections, for a total number of 2 *×* 3 *×* 4 = 24 samples.

#### 3.1.2 Gene expression profiles under sleep disruption

Gene expression profiles under sleep deprivation conditions were downloaded from GEO using the GEO ID GSE69079 [94]. The gene expression profiles were measured using an Affymetrix Mouse Genome 430 2.0 Array. Mice were in sleep or sleep-deprived conditions, and six biological replicates were used for the bound fraction. For the unbound fraction, sleep-deprived mice had two biological replicates. Sixteen mice were used in the present study.

### 3.2 Tensor decomposition based unsupervised feature extraction [10]

First, the gene expression profiles must be formed as tensors. For the anaesthesia treatment, gene expression profiles were formed as a tensor *x*_*ijkm*_ ∈ ℝ^*N ×*2*×*4*×*3^, which represents the expression of the *i*th gene at The *j*th condition (anaesthesia treatment and control) in the *k*th brain tissue in the *m*th replicate. For sleep, the gene expression profiles are formed as the tensor *x*^*′*^_*ijk*_*′*_*m*_*′*∈ ℝ^*N×*2*×*2*×*6^, which represents the expression of the *i*th gene at the *j*th condition (sleeping or waking) in the *k*th condition (bound vs. unbound) in the *m*th replicate. For the unbound condition, because only two replicates were used, each replicate was repeated three times to obtain six replicates.

We generated another tensor by multiplying the two tensors as follows:

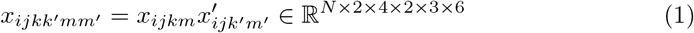

where only genes *i* common between the two profiles (sleep and anaesthesia) were considered. Gene symbols attributed to genes via bowtie2 were converted to ENTREZ GENE ID using org.Mm.eg.db package in Bioconductor. ENTREZ GENE IDs are also attributed to Affymetrix Mouse Genome 430 2.0 Array, thus we can compare ENTREZ GENE IDs between the two gene expression profiles. *j* = 1 corresponds to sleep or anaesthesia and *j* = 2 corresponds to waking or control. This enabled us to identify gene expression that correlates with the similarity between sleep/wake and anaesthesia/control, i.e. unconscious vs. conscious. *x*_*ijkk*_*′*_*mm*_*′* is scaled as

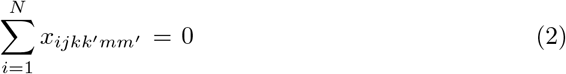

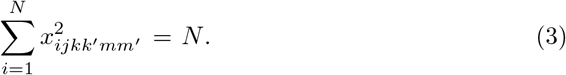

Then, HOSVD was applied to *x*_*ijkk*_*′*_*mm*_*′* to obtain

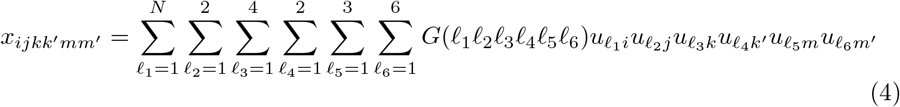

where *G* ∈ ℝ^*N ×*2*×*4*×*2*×*3*×*6^ is a core tensor that represents the contribution of individual product 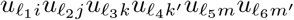 toward 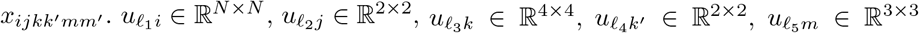, and 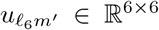, are singular value matrices and orthogonal matrices, which represent the dependence of *x*_*ijkk*_*′*_*mm*_*′* on *i, j, k, k*^*′*^, *m*, and *m*^*′*^.

After selecting 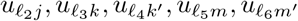 associated with desired property, we investigated *G*(*ℓ*_1_*ℓ*_2_*ℓ*_3_*ℓ*_4_*ℓ*_5_*ℓ*_6_) to see which 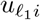 was maximally associated with the selected 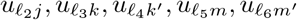. Typically, 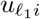 with *G*(*ℓ*_1_*ℓ*_2_*ℓ*_3_*ℓ*_4_*ℓ*_5_*ℓ*_6_), which has the largest absolute value, is selected.

Next, we selected *i*s with a larger contribution to 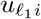. To achieve this, we attributed *P* -values to *i* by assuming that 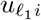 was a Gaussian

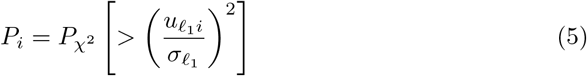

where 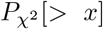 is the cumulative *χ*^2^ distribution, where the argument is larger than *x* and 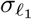 is the standard deviation. *P*_*i*_s were corrected using the BenjaminiHochberg criterion [10] to consider multiple comparison correction, and *i*s associated with adjusted *P* -values less than the threshold value (0.01) were selected.

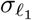 was optimized to obtain a histogram of *P*_*i*_ as flat as possible, since the histogram of *P*_*i*_ becomes uniform distribution when the null hypothesis, that is, 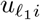, when it follows a Gaussian distribution. To identify 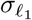 that coincides maximally with the null hypothesis, the following program was performed:

1. Set 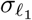.
2. Compute *P*_*i*_ with using 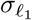.
3. Exclude *i*s associated with adjusted *P* -values less than the threshold value.
4. Re-compute *σ*_*ℓ*_ with remaining 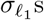.
5. Compute the histogram *h*(*P*_*i*_), of *P*_*i*_.
6. Compute the standard deviation of *h*(*P*_*i*_).
7. Update 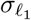.
8. Go to step 2.

This process was repeated until the standard deviation of *h*(*P*_*i*_) was minimised. This resulted in a 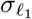 that maximally satisfied the null hypothesis because the smallest standard deviation of *h*(*P*_*i*_) corresponded to a uniform distribution, which in turn coincides with the null hypothesis as closely as possible.

### 3.3 LLM

LLMs [95] are useful for evaluating gene functions to some extent. Here, we describe how LLMs were used to retrieve significant genes from a set of selected genes.

- Gemini 2.5(experimental) + Deep Research (https://gemini.google.com/)
- perplexity Research (https://www.perplexity.ai/)
- Grok Deep Search (https://grok.com/)
- DeepSeek DeekThink(R1) + Search (https://chat.deepseek.com/)
- AI Agents for Scientific Discovery Crow (https://platform.futurehouse.org/)

The prompt used was “This list of genes has been listed as being related to sleep and anaesthesia. Please review these lists, select those that you think are relevant to both sleep and anaesthesia, and summarise them with citations to the supporting articles”, or its Japanese translation. This prompt was used to upload a list of the selected genes (one gene per line). LLMs occasionally incorrectly listed genes not in a list of 507 selected genes or 1,048 selected genes. These genes were excluded from the analysis. Among the genes selected by the LLMs, those associated with suitable references were included.

### 3.4 Enrichment analysis

Enrichment analysis was performed using Enrichr [96]. The list of symbols for the selected genes has been uploaded. *P* -values provided by Enrichr must be corrected using multiple comparison corrections and a suitable background set of genes and biological terms.

## 4 Discussion

In this study, two sets of genes, composed of 507 ad 1,048 genes, respectively, commonly associated with unconsciousness caused by sleep and anaesthesia were identified. The sets of genes included candidate genes commonly related to unconsciousness caused by sleep and anaesthesia and were identified using LLMs. In addition, using enrichment analysis, we identified various biological terms and brain subregions commonly associated with sleep- and anaesthesia-associated unconsciousness. Subsequent analysis identified genes whose expression was commonly distinct between sleep/anaesthesia and wakefulness/control. In conventional analysis, sleep and anaesthesia were studied separately, and differentially expressed genes (DEGs) were identified in both groups. Overlaps of DEGs between sleep and anaesthesia were identified. In contrast to the conventional analysis, using TD based unsupervised FE, we identified commonly distinctly expressive genes by analysing *x*_*ijkk*_*′*_*mmm*_*′*, which are the products of gene expression between sleep and anaesthesia. Only genes whose expression was expressive/suppressive between sleep or anaesthesia and the awake state can have larger/smaller values of *x*_*ijkk*_*′*_*mm*_*′*. No other methods can effectively perform this task. Despite the fully unsupervised nature of this method, TD-based unsupervised FE can be used to select a reasonable set of genes. The selected genes were found to be enriched in multiple brain subregions related to sleep and anaesthesia. Furthermore, biological terms related to sleep and anaesthesia were identified. In particular, the identification of RHO GTPases activity was significant since it had not been identified until recently.

However, contrary to expectations, gene expression analysis cannot be used to qualify the nature of unconsciousness caused by sleep and anaesthesia. Therefore, more sophisticated tools are required.

To achieve this, we generated the tensor, *x*_*ijkk*_*′*_*mm*_*′*, by multiplying two tensors, *x*_*ijkm*_ and *x*^*′*^_*ijk*_*′*_*m*_*′*. By multiplying the two tensors, *i*s that take larger or smaller values for both tensors take on larger or smaller values, respectively. In contrast, *i*s where one tensor has a larger value and the other takes has a smaller value take on medium values (i.e. neither larger nor smaller values). After standardising *x*_*ijkk*_*′*_*mm*_*′* as eqs. (2) and (3), since positive (negative) absolutely larger values are attributed to *x*_*ijkk*_*′*_*mm*_*′* with larger (smaller) values, they have tendency to have smaller *P*_*i*_ defined in eq. (5), and were selected. Ultimately, this means that the consideration of *x*_*ijkk*_*′*_*mm*_*′* enabled us to identify *i*s associated with smaller or larger values simultaneously for sleep and anaesthesia.

One might wonder whether there are justifications for assuming that 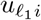 follows a Gaussian distribution. To address this, we attempted to optimise 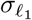 such that the histogram of *P*_*i*_ was as uniform as possible when excluding the selected *i*s. As a result, a relatively uniform distribution of the histogram of 1 − *P*_*i*_ was obtained, as shown in Figs. 3 and 4, confirming the distribution.

Although the trustworthiness of LLMs is questionable, only genes whose selection was supported by previously published studies were listed to ensure robustness in our selection.

Gene expression for sleep was obtained from astrocytes, whereas that for anaesthesia was obtained from various brain regions. Although one may wonder whether this is problematic, a comparison between these two methods has been published [9]. This indicated that a comparison between the two methods had already been conducted. In addition, because the purpose of this study was to identify commonly expressed genes, the identification of genes common between various brain subregions and astrocytes could be used to signal the robustness of our gene selection, providing evidence of trustworthiness.

We identified two sets of genes: *ℓ*_1_ = 3 and *ℓ*_1_ = 4. The proteins encoded by these genes have several distinct functions. Those selected by *ℓ*_1_ = 3 are mainly GABAergic and enriched in layer 1 of FCx, which primarily includes GABAergic neurons. In contrast, those selected by *ℓ*_1_ = 4 are mostly glutamatergic and enriched in layer 6 of PCx, which primarily includes glutamatergic neurons. The latter is closely associated with orexin. Notably, the RHO GTPases pathway was enriched in both sets of genes despite the distinct nature of the two sets of genes. Based on these results, we hypothesised that L1 GABAergic neurons in the frontal cortex and L6 glutamatergic neurons in the prefrontal cortex use RHO GTPase to differentiate between states of unconsciousness during anaesthesia and sleep. Figure 6 shows the interconnectivity between the biological terms detected by Enrichr (only those with adjusted *P* -values of less than 0.05). As expected, L1 in FCx and L6 in PCx were distinct (no shared genes). The RHO GTPase pathways share many genes, even between *ℓ*_1_ = 3 (GABAergic) and *ℓ*_1_ = 4 (glutamatergic) and are separately connected to L1 of FCx and L6 of PCx. This graph is consistent with the abovementioned hypotheses. The intersections shown in Fig. 2 may be the result of RHO GTPase pathways shared between *ℓ*_1_ = 3 (GABAergic) and *ℓ*_1_ = 4 (glutamatergic).

**Fig. 6.**
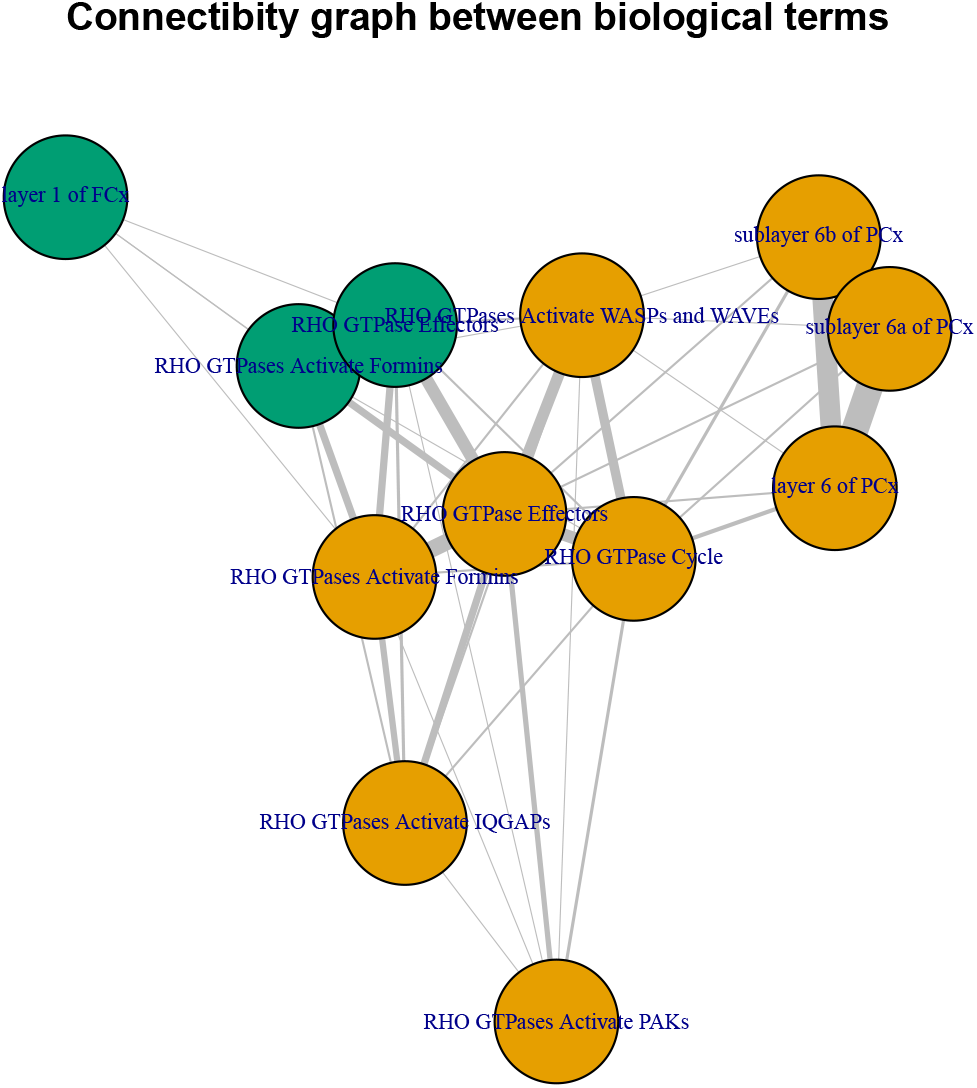
Connectivity graph between biological terms detected by Enrichr (only those with adjusted *P* -values less than 0.05). Green: *ℓ*_1_ = 3 (GABAergic), yellow: *ℓ*_1_ = 4 (glutamatergic). Edge width represents the number of genes shared between adjusted biological terms.

Although the use of a Gaussian distribution is well established [10], one might still wonder how the employment of an alternative distribution affects the selection of genes. To address this, we replaced the cumulative *χ*^2^ distribution with a cumulative exponential distribution and examined how this replacement altered the selection of genes. To the best of our knowledge, this study is the first to attempt this replacement. First, because the exponential distribution has longer tails than the Gaussian distribution, extreme values can appear easier than Gaussian. Thus, the threshold *P* - values for the exponential distribution needed to be increased. As a result, 2729 genes were found for *ℓ*_1_ = 3 and 1762 genes for *ℓ*_1_ = 4, assuming that the threshold *P* -values were 0.1. This indicates that the gene selection was not robust to distribution. Nevertheless, the enrichment analysis results were robust. For *ℓ*_1_ = 3, “Signalling by Rho GTPases, Miro GTPases and RHOBTB3” and “Signalling by Rho GTPases” were the second and the fifth highest enriched terms in “Reactome Pathways 2024”. “GABAergic synapse” was the second highest enriched in “KEGG 2021 Human”. “Layer 1 of FCx” and “layer 1 of PCx” were the top and third most enriched term in “Allen Brain Atlas down”. For *ℓ*_1_ = 4, “Signalling by Rho GTPases, Miro GTPases and RHOBTB3” and “Signalling by Rho GTPases” were the third and fourth highest enriched terms in “Reactome Pathways 2024”. “Glutamatergic synapse” was the top ranked term in terms of enrichment in “KEGG 2021 Human”. “Secondary motor area, layer 6b” was the top enriched term in “Allen Brain Atlas up”. Thus, although gene selection was not robust to different distributions, enrichment related to the key findings in this study was considered robust, independent of the assumed distribution.

Although we identified the importance of RHO GTPase, this result was fully based on a data-driven approach. This suggests that our findings should be supported by future studies. It is worth noting that the depth and duration of anaesthesia may affect the results, and that the identification of RHO GTPase is new. In this regard, the mechanism by which RHO GTPase affects unconsciousness will be the subject of a future study.

## 5 Conclusion

This study applied TD-based unsupervised FE to the gene expression of sleep and anaesthesia and identified two sets of genes that are commonly expressed distinctly between unconscious states induced by sleep or anaesthesia and wakefulness. The two sets of genes were enriched in GABAergic synapses in L1 of FCx and glutamatic synapses in L6 of PCx, and RHO GTPase pathways were shared by both sets. This suggests that GABAergic synapses in L1 of FCx and glutamatic synapses in L6 of PCx are the differentiating factors between unconsciousness induced by sleep and that induced by anaesthesia through the RHO GTPase pathways.

## Declarations

## Acknowledgements.

We thank Mr. Hiroyuki Kanaya for critically reviewing the manuscript.

## Funding

The project was funded by KAU Endowment (WAQF) at king Abdulaziz University, Jeddah, Saudi Arabia. The authors, therefore, acknowledge with thanks WAQF and the Deanship of Scientific Research (DSR) for technical and financial support.

## Conflict of interest/Competing interests

We declare that the authors have no conflicts of interest.

## Ethics approval and consent to participate

Not applicable.

## Consent for publication

Not applicable.

## Data availability

All data analysed in this study were obtained from GEO (GSE69079) and DRA010292 and are publicly available.

## Materials availability

Not applicable.

## Code availability

https://github.com/tagtag/Sleepvsanesthesia

## Author contributions

Y.-H.T. planned the study and performed the analyses. Y.-H.T. and T.T. evaluated the results and wrote and reviewed the manuscript. All authors have read and agreed to the published version of the manuscript. Conceptualization, data curation, and analysis were performed by Y.-H.T.

## Notes

### Competing Interest Statement

The authors have declared no competing interest.

### Summary of Updates

Revision due to the reviewers' comment

